# In vivo CRISPR screening identifies SAGA complex members as key regulators of hematopoiesis

**DOI:** 10.1101/2022.07.22.501030

**Authors:** Michael S. Haney, Archana Shankar, Leonid Olender, Ian Hsu, Masashi Miyauchi, Róbert Pálovics, Grace A. Meaker, Satoshi Kaito, Ola Rizq, Hwei Minn Khoo, Yavor Bozhilov, Kyomi J. Igarashi, Joydeep Bhadury, Christy Munson, Paul K. Mack, Tze-Kai Tan, Jan Rehwinkel, Atushi Iwama, Tony Wyss-Coray, Hiromitsu Nakauchi, Adam C. Wilkinson

**Affiliations:** Department of Neurology and Neurological Sciences, Stanford University School of Medicine; Stanford, CA, USA; Wu Tsai Neurosciences Institute, Stanford University; Stanford, CA, USA; Department of Pathology and Laboratory Medicine, University of Pennsylvania, Philadelphia, Pennsylvania, USA; Institute for Stem Cell Biology and Regenerative Medicine, Stanford University School of Medicine; Stanford, CA, USA; MRC Weatherall Institute of Molecular Medicine, University of Oxford; Oxford, UK; Department of Genetics, Stanford University School of Medicine, Stanford, CA, USA; Division of Stem Cell and Molecular Medicine, The Institute of Medical Science University of Tokyo, Tokyo, Japan; Department of Laboratory Medicine, Institute of Biomedicine, Sahlgrenska Academy, University of Gothenburg; Gothenburg, Sweden; ChEM-H, Stanford University; Stanford, CA, USA; Paul F. Glenn Center for the Biology of Aging, Stanford University School of Medicine; Stanford, CA, USA; Stem Cell Therapy Laboratory, Advanced Research Institute, Tokyo Medical and Dental University, Tokyo, Japan

## Abstract

The biological mechanisms that sustain the vast blood production required for healthy life remain incompletely understood. To search for novel regulators of hematopoiesis, we performed genome-wide in vivo hematopoietic stem cell (HSC)-based CRISPR knockout screens for regulators of hematopoiesis. We discovered SAGA complex members, including *Tada2b* and *Taf5l*, as key regulators of hematopoiesis. Loss of *Tada2b* or *Taf5l* strongly inhibited hematopoiesis *in vivo*, led to a buildup of immature hematopoietic cells in the bone marrow, and was associated with upregulation of interferon pathway genes. Loss of these factors also enhanced the cell outgrowth and the interferon pathway in an *in vivo* human myelodysplastic syndrome model, suggesting that loss of SAGA complex activity could contribute to hematological disease progression. In summary, this study has identified the SAGA complex as an important regulator of hematopoiesis.

## Main text

Hematopoietic stem cells (HSCs) are a rare bone marrow (BM) cell type that have the potential to self-renew or undergo lineage commitment and differentiate into one of the many hematopoietic and immune cell types^1-5^. As such, HSCs support the hematopoietic and immune systems throughout life, which play various essential roles in human health from oxygen supply to wound healing to defense against pathogens. Approximately ~10^11^ new blood cells are generated per day in humans. Hematopoietic system function generally declines with age and can result in leukopenia, anemia, myelodysplastic syndrome (MDS), and leukemia^6,7^, as well as other pathologies (e.g., atherosclerosis, osteoporosis). It is also one of the major drivers of declining immune cell function in aged individuals, predisposing aged individuals to infection and cancers^8^. Understanding the mechanisms of healthy and pathogenic hematopoiesis is therefore a key question in the field of hematology, stem cell biology, and gerontology. In particular, the regulation of gene expression is critical for healthy hematopoiesis^9,10^. However, there is still much to be explored about this process.

Tight regulation of gene expression is critical for HSC function, with perturbations known to underlie the loss of hematopoietic system homeostasis^9^. The SAGA complex is an evolutionary-conserved multiprotein complex that contains both histone acetyl-transferase and histone deubiquitinase activities^11,12^. The canonical enzymatic proteins are Kat2a and Usp22, respectively. Loss of Kat2a has previously been shown to minimally impact HSC activity^13^, perhaps due to redundancy with Kat2b, although it is important in leukemic stem cell activity^14^. Loss of Usp22 has recently been shown to induce emergency myelopoiesis^15^. Alongside these enzymatic subunits, SAGA contains several structural components including transcriptional adaptors (TADAs) including Tada2b and Tada1, and transcription-associated factors (TAFs) including Taf5l. The role of these structural components in HSC activity is poorly understood.

While hematopoietic stem and progenitor cell (HSPC)-derived hematopoiesis is a well characterized system, the application of methods to identify genetic regulators of this process through high-throughput genetic screens have been hampered by the rarity of these cells. *In vivo* pooled CRISPR/Cas9 knockout (KO) screens offer an efficient method to investigate thousands of genetic regulators for a range of biological processes^16^, and have already been applied to a number of cell lines^14,17,18^. However, the application of this system to primary HSCs is considerably more challenging because of the large numbers of cells required to perform these screens^19^. To date, *in vivo* HSC CRISPR screens have been limited to sgRNA libraries targeting 30-150 genes^20,21^. Here, by utilizing *ex vivo* HSC expansion methodology, we undertook an HSC CRISPR screen targeting ~2000 genes related to the regulation of gene expression, from which we identify and validate structural components of the SAGA complex as key regulators of hematopoiesis and MDS.

## Results

To gain new insight into the genetics of hematopoiesis, we optimized a large-scale ex vivo HSC expansion culture method^22-24^ to generate sufficient HSC numbers for CRISPR KO screens. Here we refer to the cells in these cultures as HSCs; we acknowledge that these cultures include hematopoietic progenitor cells but for simplicity we will refer to these as HSC cultures. We performed 10 pooled CRISPR KO screens targeting the entire mouse genome (10 sgRNAs/gene) in 3-week expanded primary mouse HSC cultures. To identify molecular regulators of *in vivo* hematopoiesis, we transplanted these gene KO HSCs into 65 lethally irradiated recipients (**Figure 1a**), achieving high-level donor peripheral blood (PB) chimerism (**Figure S1a**). After 10-12-weeks, BM and spleen from these mice were collected pooled, and various hematopoietic cell types purified for sgRNA sequencing: c-Kit^+^Lineage^-^ hematopoietic stem and progenitor cells (HSPCs), CD11b/Gr1^+^ myeloid cells, CD4/CD8^+^ T cells, B220^+^ B cells, and Ter119^+^ erythroid cells.

**Figure 1:**
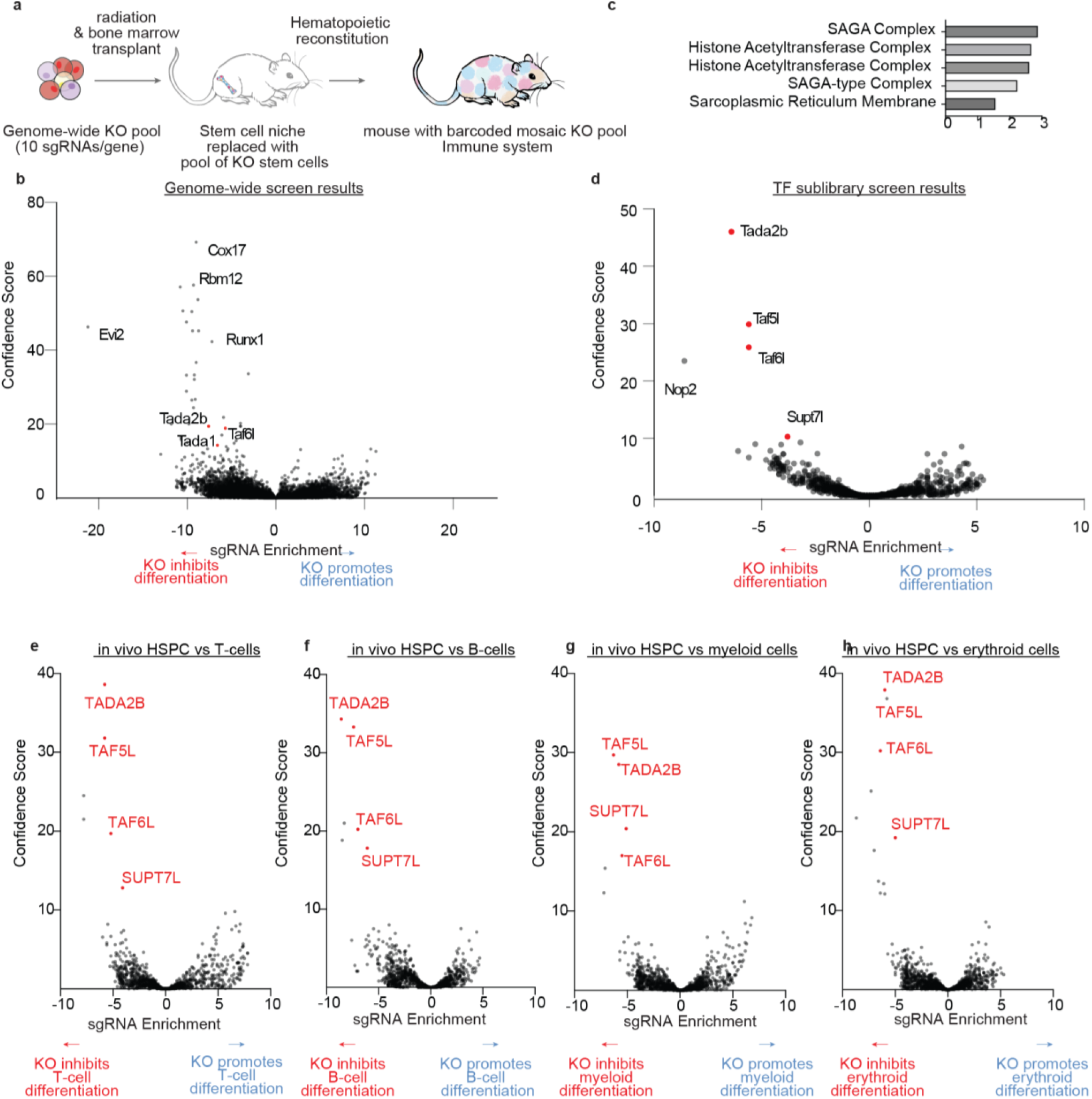
In vivo HSC CRISPR screen identifies SAGA complex members as putative regulators of hematopoiesis. (a) Schematic of the *in vivo* genome-wide (GW) CRISPR KO screen for hematopoiesis. (b) Volcano plot showing *in vivo* HSC CRISPR GW screen results comparing HSPC KOs with KO mature lineage cells. All KOs displayed the effect of KO (negative if KO enriched in HSPCs, positive if KO enriched in differentiated cells) on the x axis and confidence score on the y axis. SAGA complex members labeled in red. (c) GO enrichment of top 200 screen hits ranked by confidence score. (d) Volcano plot showing *in vivo* HSC CRISPR screen results for sublibrary of 2,000 genes involved in gene expression. All KOs displayed the effect of KO (negative if KO enriched in HSPCs, positive if KO enriched in differentiated cells) on the x axis and confidence score on the y axis. SAGA complex members labeled in red. (e-h) Volcano plot showing in vivo HSC CRISPR screen results comparing c-Kit^+^Lineage^-^ HSPC KOs with KO mature T-cells (e), mature B-cells (f), myeloid cells (g), and erythroid cells (h). All ~2,000 gene KOs displayed the effect of KO (negative if KO enriched in HSPCs, positive if KO enriched in differentiated cells) on the x axis and confidence score on the y axis.

In comparing sgRNA abundance between HSPCs and all mature lineage populations (**Figure 1b**), we identified members of the SAGA complex^11^ as being enriched within the HSPC population (**Figure 1c**). These SAGA complex members were also seen as enriched when HSPCs were compared with each individual mature lineage (**Figure S1b-e**). The role of these structural components in hematopoiesis is poorly understood, yet were the components of the SAGA complex identified in our CRISPR screen. To investigate this further, we performed a targeted 2,000 gene screen of focused on regulators of gene expression that included many of the SAGA complex members. SAGA complex members were even more pronounced as top hits that are required for normal hematopoiesis (**Figure 1d-h)**.

To validate these screen results, we repeated the HSC CRISPR KO pipeline targeting *Tada2b, Taf5l, Tada1*, or the *Rosa26* locus as a control (**Figure 2a**). Gene KOs were confirmed at the DNA and protein levels (**Figure S2a-c**). *Tada2b, Taf5l*, and *Tada1* KO cells all engrafted at much higher levels within the immature BM HSPC compartments as compared with the mature PB, while control sgRNA cells displayed similar chimerism levels throughout the hematopoietic hierarchy (**Figure 2b**). Notably, while higher levels of KO chimerism were observed in Lineage^-^ progenitor populations, chimerism within the most primitive immunophenotypic CD150^+^CD34^-^KSL LT-HSC population was similar to the PB (**Figure 2c-f**). We also observed a striking increase in the frequency of immature hematopoietic cell populations (Lineage^-^ cells and KSL HSPCs) within the KO BM cells (**Figure 2g-h**). Additionally, we confirmed that the inhibited hematopoiesis phenotype was transplantable by performing secondary transplantation assays **(Figure S2d**).

**Figure 2:**
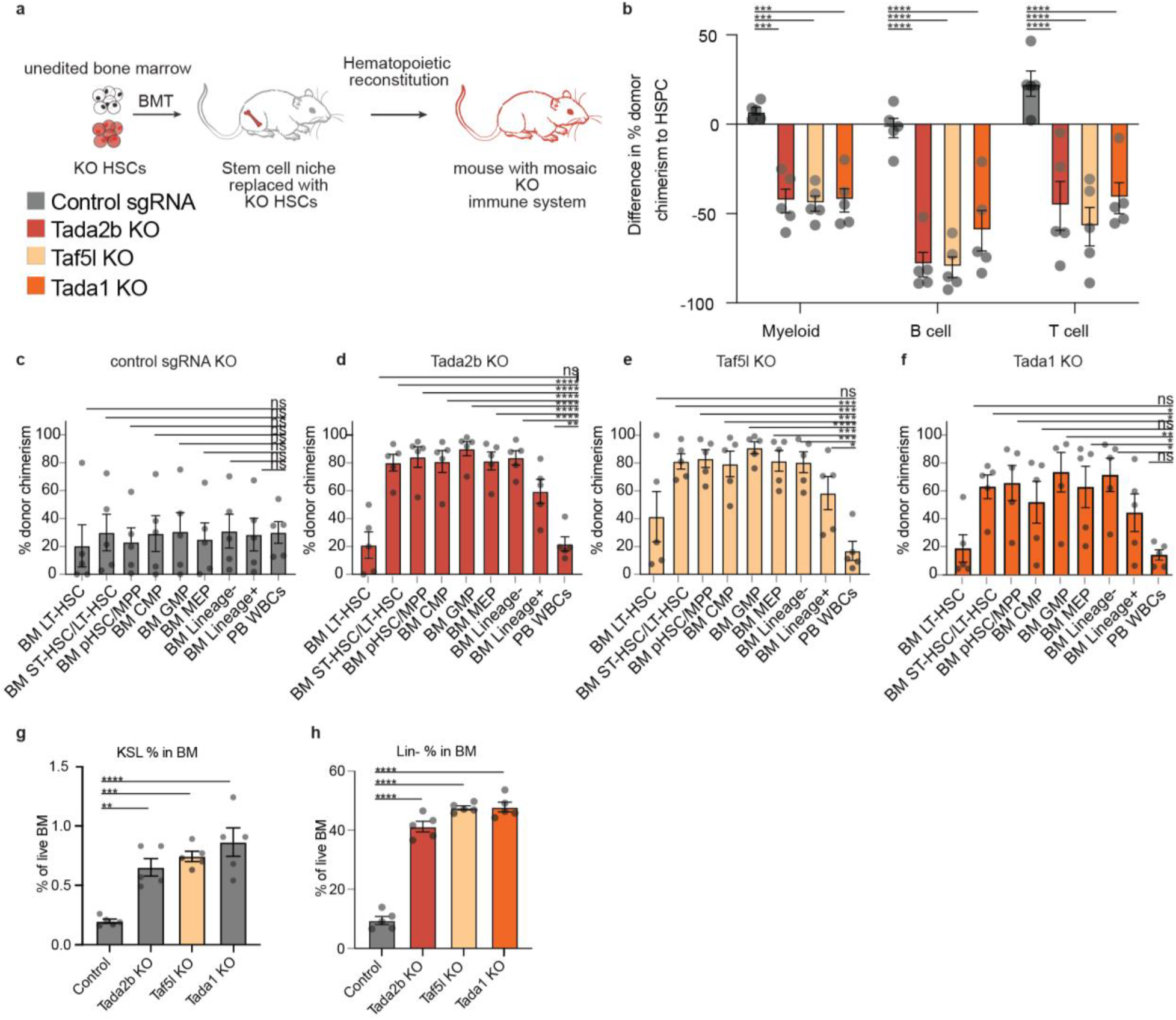
Loss of SAGA complex members inhibits normal hematopoiesis. (a) Schematic for single gene KO validations. (b) Difference in donor chimerism between the KSL HSPC compartment and myeloid, T and B cells, for transplants. Error bars represent s.e.m; P-value determined by two-way ANOVA. (c-f) Frequency of donor CD45.1^+^ chimerism in indicated hematopoietic cell populations for Rosa26 KO cells (c), Tada2b KO cells (d), Taf5l KO cells (e), and Tada1 KO cells (f). n=5 per condition. Error bars represent s.e.m; P-value determined by one-way ANOVA. (g) Frequency of c-Kit^+^Sca1^+^Lineage^-^ (KSL) cells within the CD45.1^+^ bone marrow compartment in transplant recipients at 12-weeks. Error bars represent s.e.m; P-value determined by one-way ANOVA. (h) Frequency of Lineage^-^ cells within the CD45.1^+^ bone marrow compartment in transplant recipients at 12-weeks. Error bars represent s.e.m; P-value determined by one-way ANOVA.

To confirm that these results are not confounded by extended time in *ex vivo* culture, we further validated that SAGA complex members altered the activity of functional HSCs in knock outs of *Tada2b* or *Taf5l* (using two different sgRNAs) in freshly-isolated CD150^+^CD34^-^c-Kit^+^Sca1^+^Lineage^-^ HSCs and evaluated their activity in 16-week transplantation assays. Similar to our validation experiments, donor chimerism was again lower in the PB than in the immature KSL HSPCs from the *Tada2b* and *Taf5l* KO HSCs (**Figure S2e-f**). In this transplantation assay, hematopoiesis was almost entirely donor-derived, we also observed significant reductions in white blood cell (WBC) counts in the *Tada2b*-KO and *Taf5l*-KO recipients (**Figure S2g**). Red blood cell (RBC) counts and platelet counts were not significantly reduced (**Figure S2h-i**). Together, these results confirmed SAGA complex members *Tada2b* and *Taf5l* as novel functional HSC regulators that are required for normal hematopoiesis.

We next investigated the phenotype of *Tada2b* and *Taf5l* KO in our *ex vivo* HSC cultures. Consistent with these knockouts displaying a block in differentiation, we observed an increase in the frequency of CD201^+^CD150^+^KSL population (described as the immunophenotypic HSC population *ex vivo*^25,26^) within the *Tada2b* and *Taf5l* KO cultures after 14-days (**Figure 3a**), and a corresponding loss of CD201^-^CD150^-^KSL and c-Kit^+^Sca1^-^Lineage^-^ progenitor populations (**Figure S3a-b**). This HSC expansion phenotype was also seen when quantifying total CD201^+^ live cell frequencies (**Figure S3c**). Additionally, knockdown of *TADA2B* using previously reported shRNAs^27^ within *ex vivo* human HSC cultures^28^ induced a similar increase in the frequency of immunophenotypic CD201^+^CD90^+^CD45RA^-^CD34^+^ HSC compartment (**Figure 3b**). These results provided further evidence that loss of SAGA complex components altered HSC activity and that we could investigate this phenotype *ex vivo*.

**Figure 3:**
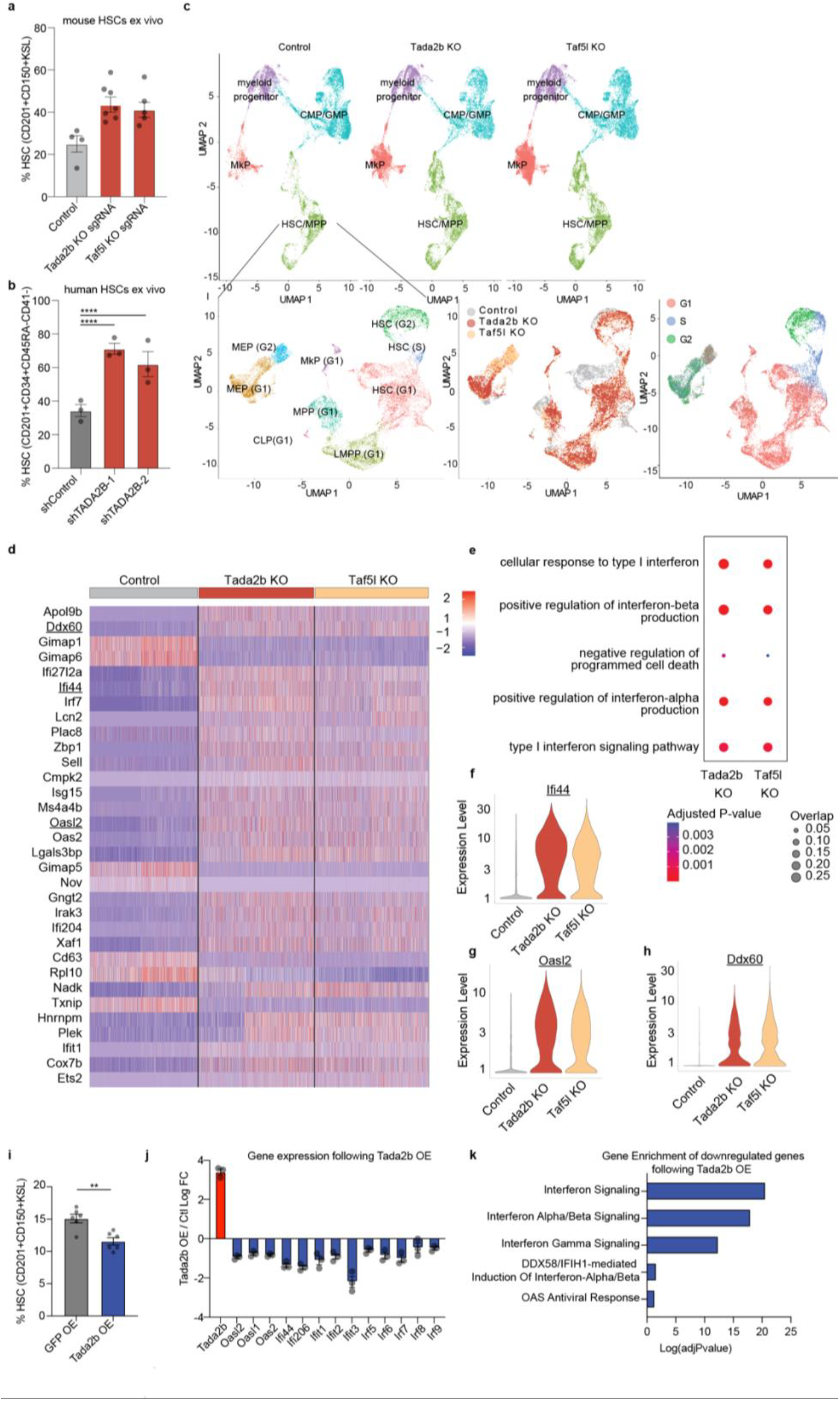
*Tada2b* or *Taf5l* regulate the interferon pathway in HSCs. (a) Percentage of phenotypic mouse HSCs (CD201^+^CD150^+^KSL) in ex vivo cultures 14-days after RNP KO of SAGA complex members. Six biological replicates with two independent sgRNAs displayed (grey: sgRNA1, black: sgRNA2). Error bars represent s.e.m; P-value determined by one-way ANOVA. (b) Percentage of phenotypic human HSCs (CD201^+^CD34^+^CD45RA^-^CD41^-^) in ex vivo cultures 14-days after shRNA knockdown of *TADA2B*. Three biological replicates displayed. Error bars represent s.e.m; P-value determined by one-way ANOVA. (c) Above, UMAP representation of single cell RNA-seq of control, *Tada2b* KO, and *Taf5l* KO mouse HSC cultures at day 14 with clusters annotated. Total of 61,324 single cells analyzed. Below, UMAP representations of the HSC/MPP cluster from a following sub-clustering with cluster annotated, cell sample origin, and cell cycle state annotated (total of 21,344 single cells analyzed). (d) Heatmap displaying the expression of the top 30 differentially expressed genes ranked by adjusted p-value between the control, *Tada2b* KO, and *Taf5l* KO cultures within the G1-phase HSC cluster from (c). Rows represent sub-sampled single-cell gene expression values. IFN response genes are highlighted in red. (e) GO enrichment analysis displaying adjusted P-value and gene-set overlap for 200 most differentially expressed genes between control and KO cells within the G1-phase HSC cluster. (f-h) Violin plots for the expression of IFN response genes *Ifi44* (E), *Oasl2* (F) and *Ddx60* (G) in control, *Tada2b* KO and *Taf5l* KO cells within the G1-phase HSC cluster. E. P-value determined by MAST. (i) Fold change of HSC (CD201^+^CD150^+^KSL) frequency in ex vivo cultures with Tada2b or GFP over-expression. Three independent replicates displayed. Error bars represent s.e.m; P-values determined by paired t-test. (j) Log fold change in expression of indicated genes in Tada2b overexpressing HSPCs. (k) GO enrichment analysis displaying adjusted P-value for downregulated genes in Tada2b overexpressing HSPCs.

Given these striking functional phenotypes from the genetic ablation of *Tada2b* and *Taf5l*, we were interested to better understand the consequences of these gene knockouts at the molecular level. We therefore performed single cell RNA-sequencing (scRNAseq) of control, *Tada2b* KO, and *Taf5l* KO HSC cultures at 14-days post-RNP electroporation. Consistent with the known heterogeneity of our HSC cultures, initial clustering analysis identified four major cell types including a population of cells with an HSC signature and several progenitor cell cultures (**Figure 3c, Figure S3d-e**). To investigate the molecular consequences specifically within the HSCs, we subclustered specifically on this population (**Figure 3c, Figure S3f**). By combining this re-clustering with cell cycle analysis, we could identify sub-clusters resembling G1-phase HSCs, S-phase HSCs, G2-phase HSCs (**Figure 3c**).

To avoid cell cycle-associated transcriptional changes, we focused on gene expression differences within the G1-HSC cluster. We observed major differences in the transcriptional programs expressed in the *Tada2b* and *Taf5l* KOs (**Figure 3d**). Gene Ontology (GO) pathway analysis identified major changes in interferon (IFN) signaling (**Figure 3e**). For example, interferon response genes *Ifi44, Ddx60*, and *Oasl2*, were significantly upregulated in the KO settings (**Figure 3f-h**). Significant upregulation of IFN-related gene sets was also observed in other progenitor clusters. The similarity of these DEGs in the *Tada2b* and *Taf5l* KOs also implicate loss of SAGA complex activity as a common mechanism underlying these transcriptional changes. Corresponding with this gene signature, we observed upregulation of known IFN response genes Sca1 (Ly6e) and MHC-I (**Figure S3g-h**).

We were interested in whether similar gene expression changes were also induced in HSPCs *in vivo*. We therefore performed bulk RNA-seq on control, *Tada2b*-KO, and *Taf5l*-KO KSL HSPCs from the BM of our transplantation recipients. We also observed increases in IFN response genes in this context, suggesting this phenotype was not an artifact of the HSC culture system but was tightly linked to the loss of these SAGA components (**Figure S4a-c**). However, within the *in vivo* setting, we could not observe changes in Sca1 (nor MHC-I) by flow cytometry (**Figure S4d-e**).

To test whether SAGA complex members are not only necessary for restraining IFN signaling but sufficient for suppressing IFN genes, we overexpressed *Tada2b* in *ex vivo* expanded HSCs. We observed a decrease in CD201^+^CD150^+^KSL HSC percentages in *ex vivo* HSC cultures upon *Tada2b* overexpression (**Figure 3i**), the opposite of the KO phenotype. *Tada2b* overexpression resulted in a suppression of several IFN-related genes and was the top GO category in the set of significantly downregulated genes (**Figure 3j-k, Figure S4f**), indicating that Tada2b expression is both necessary and sufficient to regulated IFN gene expression in HSPCs.

To understand where SAGA was influencing the intrinsic IFN pathway, we initially assessed inhibition of the upstream IFN pathway regulator Tbk1. Inhibition led to ~50% reduction in the expression of the ISG Sca1 (**Figure 4a**). Similar effects were seen in *Tada2b*-KO and control backgrounds, suggesting that Tbk1 activity was not unique to the *Tada2b*-KO setting. We next assessed whether IFN secretion (and its signaling) could be contributing to the SAGA complex KO phenotype by performing an IFN secretion assay. We detected higher IFN levels in *Tada2b* KO cultures as compared with control cultures (**Figure 4b**).

**Figure 4:**
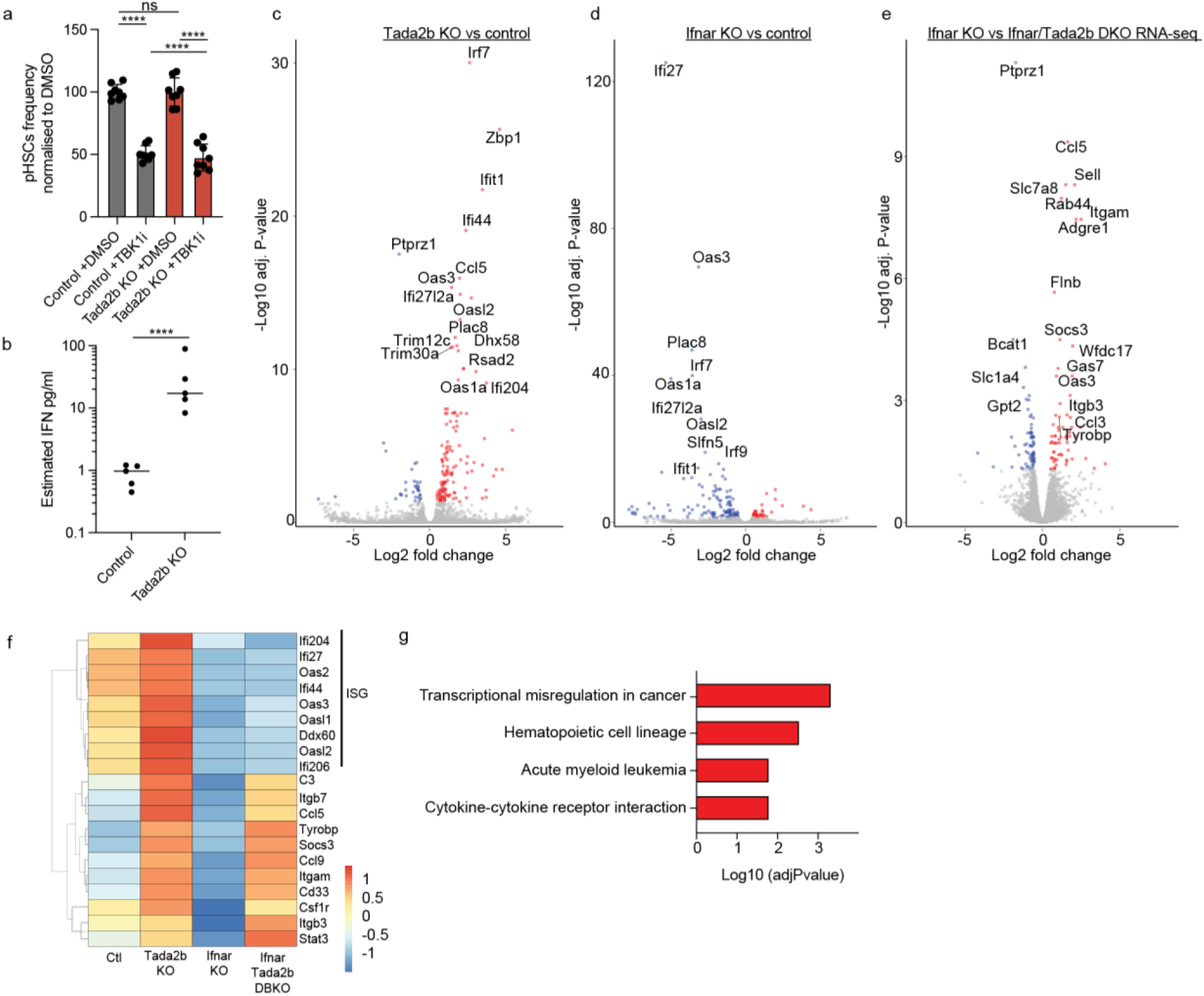
SAGA complex members have IFN dependent and independent effects on HSCs. (a) Sca1 MFI in HSC cultures following a 7-day treatment with the TBK1 inhibitor GSK-8612. P-values determined by one-way ANOVA. (b) Estimated concentration of interferons in the supernatant of 14-day *Tada2b* KO or control KO HSC cultures. Supernatant collected 48 hours after the last media change. P-values determined by t-test. (c) Volcano plot comparing gene expression between control CD201^+^CD150^+^KSL HSCs and *Tada2b-*KO CD201^+^CD150^+^KSL HSCs, at 14-days culture after RNP electroporation. (d) Volcano plot comparing gene expression between control CD201^+^CD150^+^KSL HSCs and *Ifnar-*KO CD201^+^CD150^+^KSL HSCs, at 14-days culture after RNP electroporation. (e) Volcano plot comparing gene expression between *Ifnar1-*KO CD201^+^CD150^+^KSL HSCs and *Ifnar1/Tada2b* double-KO CD201^+^CD150^+^KSL HSCs, at 14-days culture after RNP electroporation. (f) Heatmap displaying expression of indicated genes within indicated HSC cultures. (g) GO enrichment analysis displaying adjusted P-value for upregulated genes indicated in (e).

To determine the contribution of secreted IFN(alpha/beta) on the SAGA complex KO-associated gene expression changes, we knocked out *Tada2b* in *Interferon alpha/beta receptor alpha chain* (*Ifnar1*)-deficient HSCs and performed bulk RNA-seq. The upregulation of IFN response genes was lost when *Tada2b* was knocked out in the *Ifnar1*-KO background (**Figure 4c-e**), implicating IFNs as mediators of this gene expression pattern. However, another set of genes were similarly upregulated following loss of *Tada2b* in both the WT and *Ifnar1*-KO backgrounds (**Figure 4f-g)**. Additionally, the increased CD201^+^CD150^+^KSL frequency phenotype remained in the SAGA complex KOs in the *Ifnar1*-deficient background (**Figure S4g**). These results implicate IFN signaling as one of several molecular pathways regulated by SAGA in HSCs, in line with the multifaceted functions of the SAGA complex^11^.

The dramatic block in hematopoiesis was reminiscent of the failure seen in hematological diseases such as MDS. We therefore set out to evaluate whether loss of SAGA complex influenced a MDS using the human MDS-L cell line model^29^. We knocked out SAGA complex members *TADA2B, TAF5L, TADA1*, or the *AAVS1* control locus using CRISPR (**Figure S4h**). We then performed a competitive transplantation assay into immunodeficient mice expressing human IL-3 and GM-CSF (**Figure 5a**). We observed a striking competitive advantage by the SAGA component KO MDS cells as compared with the control cells, with significantly higher chimerism in both the BM and spleen (**Figure 5b-c, Figure S4i**). Additionally, we observed larger spleen weights in *TADA1* and *TAF5L* KO recipients (**Figure S4j**). However, there were no overt signs of leukemic transformation (**Figure S4k**) or recipient mortality.

**Figure 5:**
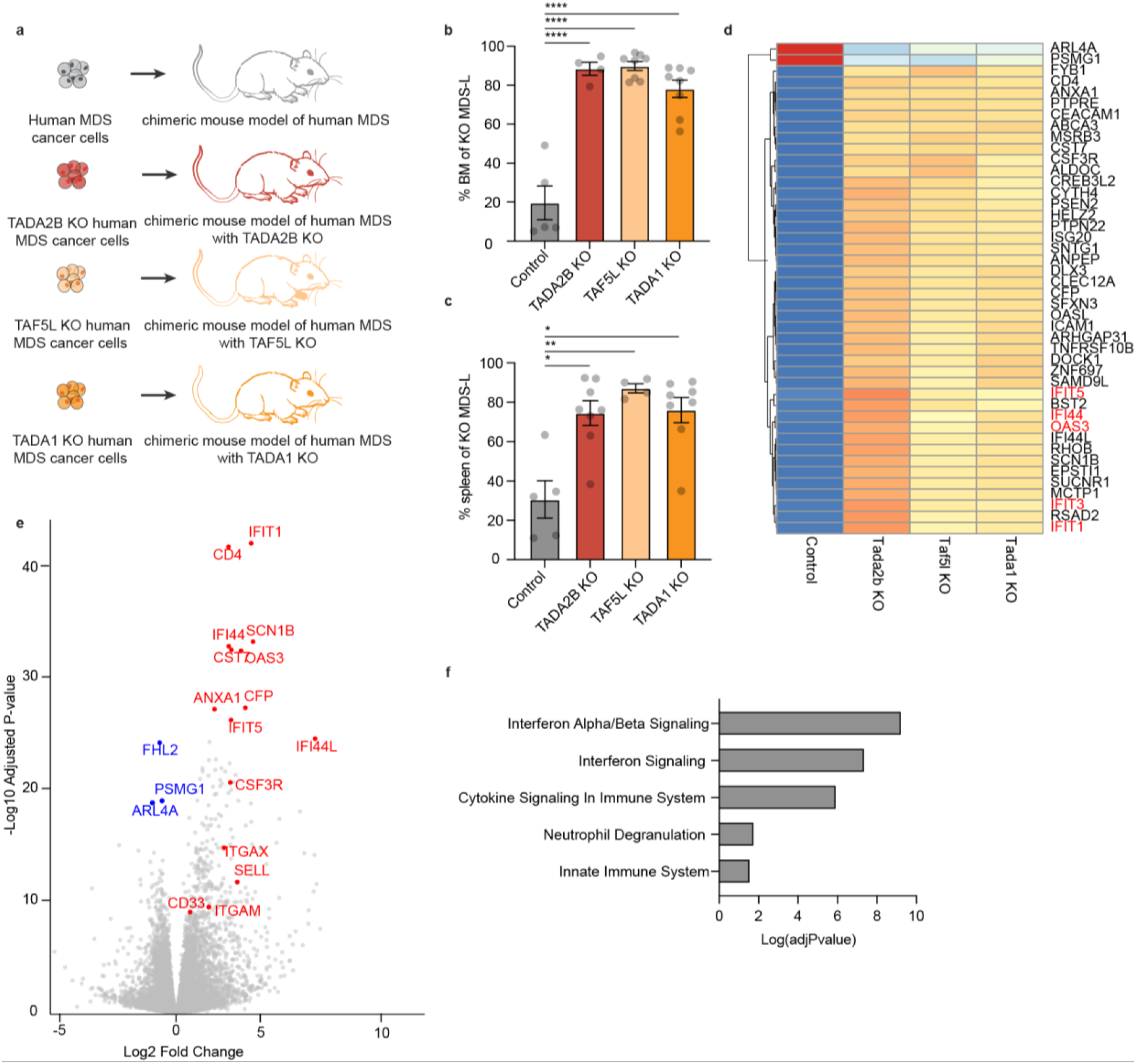
SAGA complex members regulate cell outgrowth in myelodysplastic syndrome. (a) Human MDS chimeric mouse model schematic. (b) Frequency of KO MDS cells in bone marrow. P-value determined by one-way ANOVA. Error bars represent s.e.m. ****P<.0005. (c) Frequency of KO MDS cells in spleen. P-value determined by one-way ANOVA. Error bars represent s.e.m. **P<.005, *P<.05. (d) Heat map comparing gene expression between *in vivo* MDS control cells with in vivo MDS *TADA2B* KO, *TAF5L* KO and *TADA1* KO cells. (e) Volcano plot comparing gene expression between *in vivo* MDS control cells with in vivo MDS *TADA2B* KO cells. (f) GO enrichment of upregulated genes in MDS *TADA2B* KO cells as compared with control cells.

To see if the same transcriptomic signature were present in the human SAGA complex KOs as in the mouse setting, we performed RNA-seq on the 12-week BM engrafted human SAGA complex KOs cells as well as controls. We again observed a robust upregulation of IFN response genes upon SAGA complex member KOs (**Figure 5d-e**). In addition to these IFN genes, we also observe an upregulation of AML and myelodysplastic-linked genes, overlapping with those found in the *Ifnar1/Tada2b* double KO HSCs (**Figure S4l**). Together, these results suggest the SAGA complex has various regulatory activities that are conserved between mouse and human.

## Discussion

In this study, we identified and validated structural components of SAGA complex members, *Tada2b, Taf5l*, and *Tada1*, as novel regulators of hematopoiesis. Loss of these factors led to buildup of immature HSPCs in the bone marrow, and a loss of mature blood cells in the peripheral blood following transplantation. At the transcriptional level, this was associated with an upregulation in IFN response genes in HSCs *ex vivo* and *in vivo*, likely due to the increased secretion of IFN. However, loss of IFNa/b sensitivity did not entirely reverse the SAGA KO HSC phenotype, suggesting additional mechanisms of SAGA in HSCs that warrant further investigation.

We also identified a role for these SAGA complex members in a human MDS model, where loss of these SAGA complex members enhanced outgrowth *in vivo*. Upregulation of IFN response genes was also seen in this human cell context, suggesting a conserved molecular pathway in mouse and human. However, it remains to be determined whether SAGA directly or indirectly induces this IFN response. Given the functional studies of the SAGA complex members we describe here, we hypothesize that loss of SAGA complex activity might be contributing to age-related dysfunction of hematopoiesis. Our findings also suggest that these structural components warrant study in other adult stem cell systems.

This study also highlights the potential for combining genetic screens with novel HSC expansion approaches. The applications of this HSC CRISPR screening approach are abundant; from identifying regulators of self-renewal and hematopoiesis, to investigating mechanisms in the full range of biological functions of mature HSC-derived immune cells or searching for synthetic lethality gene interactions in pre-malignant HSCs.

## Materials and Methods

### Mice

All animal experiments were approved by the Administrative Panel on Laboratory Animal Care at Stanford University, the UK Home Office, or the Animal Care and Use Committee of the Institute of Medical Science University of Tokyo. C57BL/6-Rosa26^CAG-Cas9^ (B6J.129(Cg)-Gt(ROSA)26Sor^tm1.1(CAG-cas9*,-EGFP)Fezh/J^; 026179) and C57BL/6-CD45.1 (PepboyJ; 002014) were purchased from The Jackson Laboratory. C57BL/6-CD45.2 mice purchased from The Jackson Laboratory (000664) or bred at the University of Oxford. C57BL/6-CD45.1/CD45.2 F1 mice were bred from C57BL/6-CD45.1 and C57BL/6-CD45.2 at Stanford University or the University of Oxford. NOD.Cg-Prkdc^scid^ Il2rg^tm1Sug^ Tg(SRa-IL3, CSF2)7-2/Jic (NOG-EXL) mice expressing human IL-3 and GM-CSF (NOG/IL-3/GM-CSF) were purchased from In-Vivo Science Inc. *Infar1*-KO (B6.bkg-Ifnar1^tm1Agt/Mmjax^) mice were provided by Caetano Reis e Sousa. All mice were 8-12 weeks at the experiment start point.

### Ex vivo mouse HSC cultures

Mouse HSC cultures were initiated from c-Kit^+^ bone marrow^30^ from C57BL/6-Rosa26^CAG-Cas9^ mice. Tibia, femur, pelvis, and vertebrae were collected and crushed to release bone marrow cells. Bone marrow was then stained with c-Kit-microbeads (Miltenyi) and c-Kit^+^ cells enriched via MACS column (Miltenyi) according to the manufacturer’s protocol. Enriched c-Kit^+^ bone marrow cells were cultured in HSC expansion media with complete media changes at 5% CO_2_ and 5% O_2_ with complete media changes every 2-3 days. HSC expansion media composition^24,31^: Ham’s F12 media supplemented with 1X penicillin-streptamycin-glutamine (PSG; Gibco), 1X HEPES, 1X insulin-transferrin-selenium-ethanolamide, 1 mg/ml polyvinyl alcohol (Sigma), 100 ng/ml recombinant mouse thrombopoietin (Peprotech), and 10 ng/ml recombinant mouse stem cell factor (Peprotech). For validation experiments, HSC cultures were initiated with c-Kit^+^ bone marrow from C57BL/6-CD45.1, C57BL/6-CD45.2, or *Ifnar1*-KO mice, as indicated in the text. Where indicated, HSC cultures were supplemented with the TBK1 inhibitor GSK-8612 at 5 uM or DMSO.

### Ex vivo and in vivo CRISPR screen

After 3-week *ex vivo* expansion of c-Kit^+^ bone marrow HSCs, ~50 million cells were transduced overnight with lentivirus generated from the Bassik mouse CRISPR knockout sub-libraries^32^ (a kind gift from the Bassik lab; Addgene 1000000121-1000000130) generated using helper plasmids pCMV-VSV-G-RSV-Rev and pMDLg/pRRE. To reduce multiple integrations per cell, we used a transduction efficiency of 20-30%. Two days later, transduced HSCs were selected for using puromycin for 72 hours. HSCs were recovered and expanded in culture for another 9 days. At day-14 post-transduction, HSCs were transplanted into lethally-irradiated (10Gy) C57BL/6-CD45.1 recipients (1-2 million HSCs per recipient). After 10-12-weeks, whole bone marrow and spleen were harvested. From the whole bone marrow, c-Kit^+^Sca1^+^Lineage^-^ or c-Kit^+^Lineage^-^ cells were isolated by FACS. The remaining bone marrow and spleen cells were pooled and CD45R^+^ B cells, CD4/CD8^+^ T cells, Mac1/Gr1^+^ myeloid cells, and Ter119^+^ erythroid cells were isolated via MACS columns.

### Screen analysis

Genomic DNA was extracted for all screen populations separately according to the protocol included with QIAGEN Blood Mini or Midi Kit. Using known universal sequences present in the lentivirally incorporated DNA, sgRNA sequences were amplified and prepared for sequencing by two sequential PCR reactions as previously described^32^. Products were sequenced using an Illumina Nextseq to monitor library composition (20–40 million reads per sample). Trimmed sequences were aligned to libraries using Bowtie, with zero mismatches tolerated and all alignments from multi-mapped reads included. Guide composition and comparisons across each cellular fraction were analyzed using CasTLE version 1.0^32^. Briefly, enrichment of individual sgRNAs was then calculated as a median-normalized log ratio of the fraction of counts. For each gene, a maximum likelihood estimator was used to identify the most likely effect size and associated log-likelihood ratio (confidence score) by comparing the distribution of gene-targeting guides to a background of nontargeting and safe-targeting guides. P-values were then estimated by permuting gene-targeting guides, and hits were called using FDR thresholds calculated via the Benjamini-Hochberg procedure.

### Ex vivo validation assays

To generate *Taf5l* or *Tada2b* KO HSCs, 2×10^5^ HSCs derived from two-week C57BL/6 c-Kit^+^ bone marrow (as described in the ***Ex vivo* HSC culture** section) were electroporated with 6ug recombinant HiFi Cas9 (IDT) pre-complexed with 3.2ug synthetic sgRNA (Synthego; sequences listed in the **Methods Table**) using a Lonza 4D-Nucleofector using program EO100 in primary P3 solution and then returned to HSC expansion media^33^. Electroporation of Cas9 only or *Rosa26*-targeting RNP were used as controls. At indicated timepoints, cells were collected for KO frequencies or flow cytometry analysis. To determine KO frequencies, gDNA was isolated via QuikExtract for PCR (primer sequences listed in the **Methods Table**) and Sanger sequencing with KO frequencies determined using ICE software (Synthego). For flow cytometric analysis, cells were antibody stained (CD201-APC, CD150-PE/Cy7, cKit-BV421, Sca1-PE, Gr1-APC/eFluor780, Ter119-APC/eFluor780, CD4-APC/eFluor780, CD8-APC/eFluor780, CD45R-APC/eFluor780, CD127-APC/eFluor780) for 30 minutes at 4°C, washed and then analyzed using a LSRFortessa (BD) with propidium iodide (Biolegend) as a live/dead cell stain. Alternatively, lentivirus carrying Tada2b-2A-GFP cDNA, or GFP cDNA were generated and transduced into 3-week old Cas9-expressing (or wild-type) C57BL/6 HSC cultures, as described in the **Ex vivo and in vivo CRISPR screen** section. For interferon studies, two-week C57BL/6 c-Kit^+^ bone marrow cells were cultured in media with recombinant mouse interferon alpha (Biolegend) as indicated with media changes every 2-3 days. Flow cytometric analysis was then performed following antibody staining (CD201-APC, CD150-PE/Cy7 cKit-BV421, Sca1-PE, Gr1-FITC, Ter119-FITC, CD4-FITC, CD8-FITC, CD45R-FITC, CD127-FITC) for 30 minutes at 4°C, washed and then analyzed using a LSRFortessa (BD) using propidium iodide (Biolegend) as a live/dead cell stain. All flow cytometric data analysis was performed using FlowJo and statistical analysis performed using Prism.

### Western blotting

Protein lysates extracted from control and KO HSCs were run on a 4-12% Bis-Tris gel and blotted using antibodies against Tada2b (St. John’s Laboratory, STJ194361-200), Taf5l (Proteintech, 19274-AP), Tada1 (Proteintech, 20337-1-AP), Gapdh (Bethyl Laboratories, A300-641A), or histone H3 (Abcam, ab1791). Following secondary staining with peroxidase-conjugated goat anti-rabbit IgG (Sigma Aldrich, A6667), blots were imaged using a ChemiBlot Imager and the Precision Plus Protein Kaleidoscope kit (Biorad 1610375).

### Transplantation assays using RNP-KO HSCs

CD150^+^CD34^-^Kit^+^Sca1^+^Lineage^-^ HSCs were isolated using a FACSAriaII (BD) from C57BL/6-CD45.1 bone marrow and electroporated with either Cas9/sgRNA targeting *Tada2b, Taf5l*, or *Rosa26* (same protocol as **Ex vivo validation assays** section) and then expanded for 7-days before transplantation into lethally-irradiated (10 Gy) C57BL/6-CD45.2 recipient mice (500 HSC equivalent per recipient) alongside 1 million whole bone marrow competitor cells isolated from C57BL/6-CD45.1/CD45.2 F1 mice. Peripheral blood samples were analyzed using a Horiba Micros ESV 60 complete blood counter or by flow cytometry. For flow cytometry, peripheral blood leukocytes were stained with antibodies (CD45.1-PE/Cy7, CD45.2-BV421, Mac1-PE, Gr1-PE, CD45R-APC/Cy7, CD4-APC, CD8-APC) following red blood cell lysis. Samples were then run on an FACSAriaII or LSRFortessa using propidium iodide as a live/dead cell stain, with CD45.1^+^ cells sorted for KO analysis. Data analysis was performed using FlowJo. At the endpoint, bone marrow and spleen were collected and fixed (4% paraformaldehyde for 48 hours) for hematoxylin and eosin staining (performed by the Stanford Animal Histology Service). Bone marrow chimerism was also analyzed by sequential antibody staining with a lineage cocktail (Gr1-biotin, Ter119-biotin, CD4-biotin, CD8-biotin, CD45R-biotin, CD127-biotin) followed by Streptavidin-APC/eFluor780, CD34-FITC, c-Kit-APC, Sca1-PE, CD45.1-PE/Cy7, CD45.2-BV421). Samples were then run on an FACSAriaII or LSRFortessa using propidium iodide as a live/dead cell stain, with CD45.1^+^c-Kit^+^Lineage^-^ cells sorted for KO analysis. All flow cytometric data analysis was performed using FlowJo and statistical analysis performed using Prism. Antibody information is listed in the **Methods Table**.

### Transplantation assays using RNP-KO HSPCs

C57BL/6-CD45.1 c-Kit^+^ bone marrow from were expanded for 14-days and then electroporated with either Cas9/sgRNA targeting *Tada2b, Taf5l, Tada1, Runx1* or *Rosa26* (same protocol as **Ex vivo validation assays** section). After a second 14-day culture, cells were transplanted into lethally-irradiated C57BL/6-CD45.2 recipient mice (5,000 CD201^+^CD150^+^c-Kit^+^Sca1^+^Lin^-^ cells per recipient) alongside 1 million whole bone marrow competitor cells isolated from C57BL/6-CD45.1/CD45.2 F1 mice. Recipient mice were analyzed as described in the **Transplantation assays using RNP-KO HSCs** section. For secondary transplantation assays, 5 million whole bone marrow cells from primary recipients were transplanted into lethally-irradiated recipient mice and analyzed as above.

### Interferon secretion assay

NIH/3T3 cells were incubated with HSC culture supernatant (48 hours after the last media change) or with recombinant IFNa (dilution series from 100 ng/ml to 10 pg/ml; Biolegend, 752802) for 3.5 hours and then collected for RNA extraction using the Quick-RNA microprep kit (Zymo Research, R1055). Following cDNA synthesis using the SSIV Reverse Transcriptase (ThermoFisher, 18090050), Taqman qPCR was run to quantify Ifit1 (ThermoFisher, Mm00515153_m1) and Gapdh (ThermoFisher, Mm99999915_g1) expression.

### Human shRNA knockdown assay

Human primary cell experiments were approved by the University of Oxford’s research ethics committee (OxTREC-574-23). Four-day cultured human umbilical cord blood CD34^+^ HSPCs (originally purchased from NHSBT) were transduced with lentivirus carrying control or *TADA2B* targeting shRNAs (reported previously^27^) and were then cultured for 14 days. HSPCs were cultured in 3A media conditions^28^: IMDM supplemented with Butzyamide (Cellaid), 740 Y-P (MedChemExpress), UM729 (StemCellTechnologies), and FLT3L (Peprotech). HSPCs were then stained with CD34-APC/Cy7, CD41-APC, CD45RA-BV785, and CD201-PE and analyzed by flow cytometry using PI as a live/dead cell stain.

## Human MDS-L studies

Human mScarlet-expressing MDS-L cells^29^ (a kind gift from Tohyama laboratory, Kawasaki Medical School) were generated previously^34^ and maintained in RMP1640 supplemented with 10% fetal bovine serum, 1X PSG, and 10ng/ml human IL-3 (Peprotech). MDS-L/mScarlet/Cas9 cells were initially generated using pLentiCas9-Blast lentivirus (Addgene 52962). MDS-L/mScarlet/Cas9 cells were then transduced and Puromycin-selected with indicated pMCB306 lentivirus expressing sgRNA and GFP (a kind gift from the Bassik lab; Addgene 89360). After confirming high KO efficiency, a 1:1 mixture of MDS-L/Cas9 cells and MDS-L/mScarlet/Cas9/sgRNA cells were then transplanted into NOG mice expressing human IL-3 and GM-CSF. After 12-weeks, bone marrow and spleen were harvested from recipient mice for flow cytometric analysis.

### Single-cell RNAseq analysis

At day-14 post-electroporation, control, *Tada2b* KO, or *Taf5l* KO HSC cultures were collected for RNA-seq analysis using a Gene Expression kit v3 (10x Genomics) according to the manufacturer’s protocol with libraries sequenced on a NovaSeq 6000 S4 (Illumina). Raw gene counts were obtained by aligning reads to the mouse genome using CellRanger software (v.4.0.0) (10x Genomics). For quality control, cells with unique feature counts over 2,500 or less than 200 were removed as well as cells that have >5% mitochondrial counts. Ambient cell free mRNA contamination was removed using SoupX for each individual sample. Multiplets were filtered out using DoubletFinder. The core statistical parameters of DoubletFinder (nExp and pK) used to build artificial doublets for true doublet classification were determined automatically using recommended settings. The SCTransform-based integration workflow of Seurat was used to align data, using default settings. In brief, the integration workflow searches for common gene modules (anchors) in cells with similar transcriptomes. Individual samples after undergoing quality control are integrated in a stepwise fashion, using cellular sequencing depth as a covariate to mitigate technical artifacts. After combining the samples into a single dataset or Seurat object, genes were projected into principal component space using the principal component analysis (RunPCA). The first 100 dimensions were used as inputs into the FindNeighbours, FindClusters and RunUMAP functions of Seurat. In brief, a shared-nearest-neighbor graph was constructed based on the Euclidean distance metric in principal component space, and cells were clustered using the Louvain method. RunUMAP functions with default settings was used to calculate 2D UMAP coordinates and search for distinct cell populations. Distinct cell populations were annotated based on conical marker genes. Monocle3 (v.0.2.1.) was used to generate the pseudotime trajectory analysis of HSCs. Cells were re-clustered as described in and used as input into Monocle to infer cluster and lineage relationships within a given cell type. Specifically, UMAP embeddings and cell subclusters generated from Seurat were converted to a cell_data_set object using SeuratWrappers (v.0.2.0) and then used as input to perform trajectory graph learning and pseudotime measurement through reversed graph embedding with Monocle. Cell cycle analysis was performed in Seurat using a list of cell cycle markers from Tirosh et al^35^. Differential gene expression was performed in Seurat using the MAST package under default conditions.

## Acknowledgments

We thank the Stanford Stem Cell Institute FACS Core and WIMM Flow Cytometry Core for flow cytometry access, the Stanford Animal Histology Service for histology, the CZ-Biohub for performing the next generation sequencing, and the WIMM Genome Engineering Core for plasmid cloning. We thank Amanda Ghassaei for assistance with the interactive webapp and figures. This research was supported by an IMSUT Collaborative Research Project. A.C.W. acknowledges support from the Kay Kendall Leukaemia Fund, the European Hematology Association, the MRC, the NIH (K99HL150218), the Leukemia and Lymphoma Society (3385-19), and the Edward P. Evans Foundation. H.N. was supported by the NIH (R01DK116944; R01HL147124), the Ludwig Foundation, and the Japan Society of the Promotion of Science. J.B. was supported by an international postdoc grant (2017-0034) from the Swedish Research Council and the Assar Gabrielsson Foundation, Sweden. M.S.H acknowledges support from the Buck Institute and NIA (T32AG000266). T.W.C. acknowledges support from the NIH (AG072255).

## Authorship contributions

MH, AS, LO, IH, MM, RP, GAM, SK, OR, HMK, YB, KJI, JB, CM, PKM, TKT, JR, AI, and ACW performed experiments and analyzed data. MH, AS, LO, HN, TWC, and ACW wrote the manuscript. All authors reviewed and edited the manuscript.

## Data availability

The datasets and materials generated and analyzed during the current study are available from the corresponding author upon request. We have made several CRISPR screening datasets publicly accessible via an interactive web app: www.hematopoiesiscrisprscreens.com.

