## Supplementary figures for "In vivo CRISPR screening identifies SAGA complex members as key regulators of hematopoiesis"

**Figure S1: In vivo GW and GE-CRISPR KO screens**

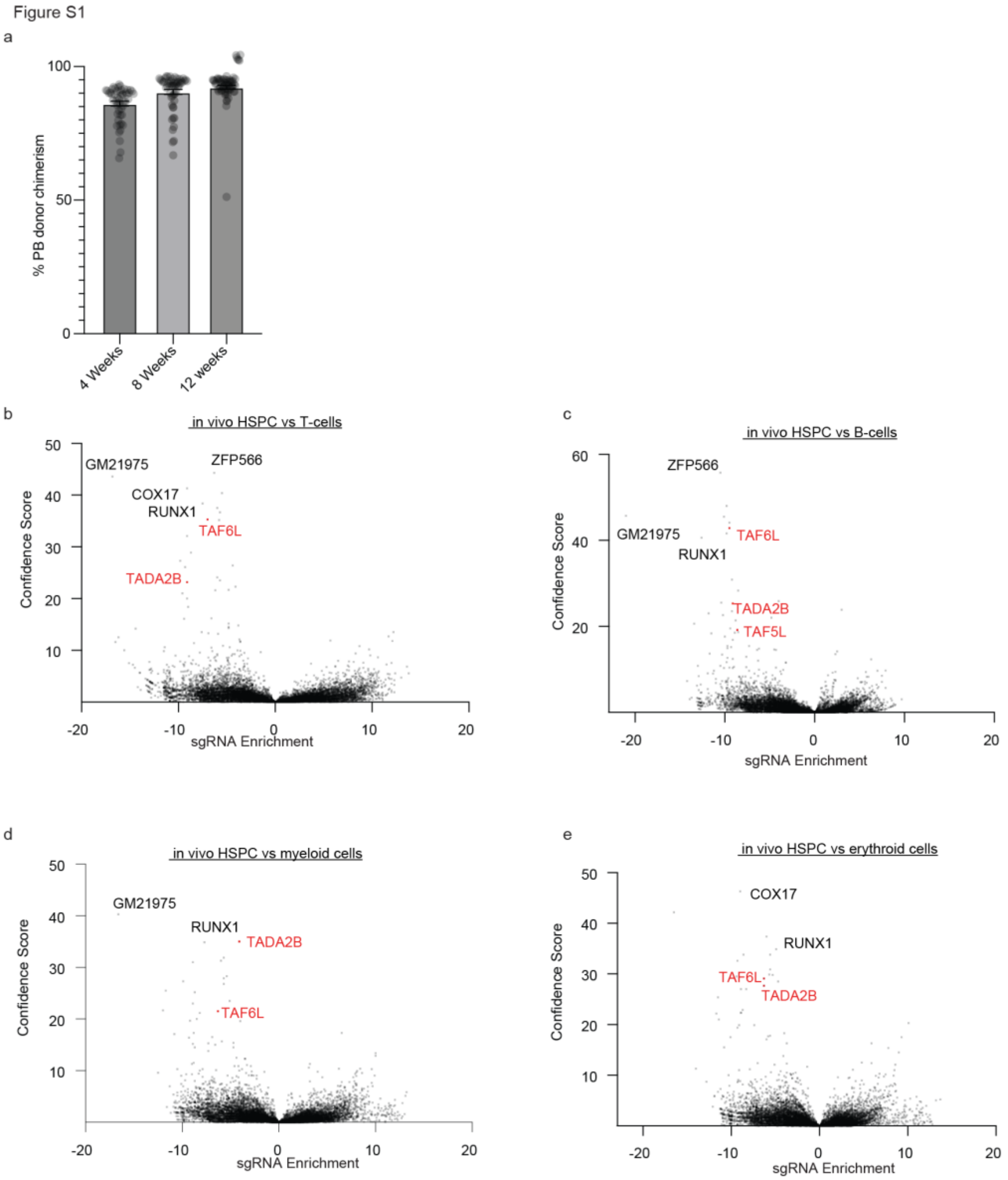

(a) Peripheral blood chimerism of GW CRISPR library cells across recipient mice.

(b-e) Volcano plot showing in vivo HSC CRISPR screen results comparing c-Kit$^{+}$Lineage$^{-}$ HSPC KOs with KO T-cells (b), B-cells (c), myeloid cells (d), and erythroid cells (e). All gene KOs displayed the effect of KO (negative if KO enriched in HSPCs, positive if KO enriched in differentiated cells) on the x axis and confidence score on the y axis.

**Figure S2: Analysis of SAGA component KO HSCs**

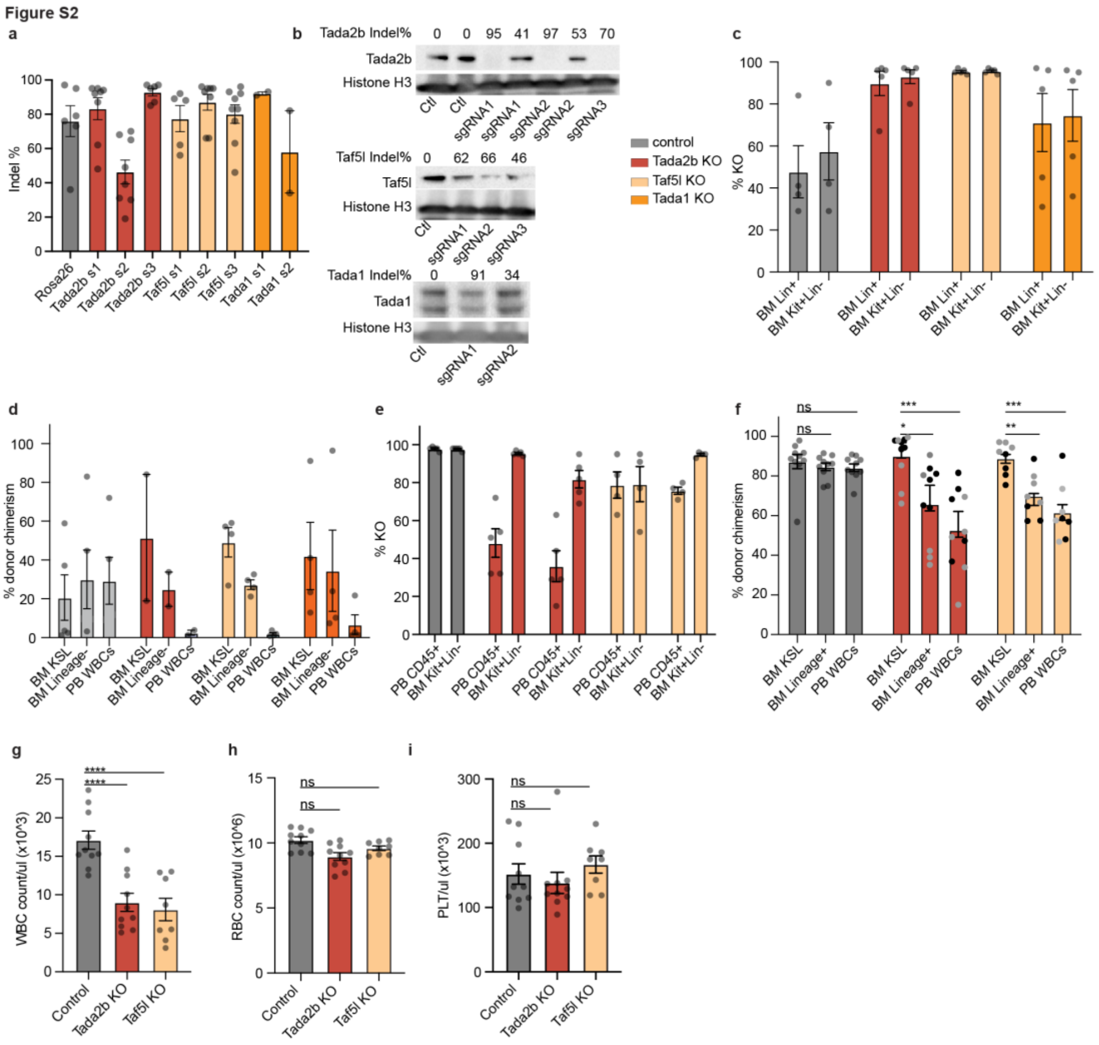

(a) INDEL frequencies with sgRNAs used in this study.

(b) Western blots showing Tada2b, Taf5l, and Tada1 protein levels in KO cells. Percentage INDELS for each sample indicated above.

(c) Gene KO frequencies in bone marrow cells 12-weeks after transplantation described in Figure 1f.

(d) Donor chimerism within bone marrow and peripheral blood within secondary recipients at 16-weeks post-transplantation for transplants described in Figure 1e.

(e) Gene KO frequencies in indicated bone marrow populations in primary recipients 16-weeks post-transplantation of freshly-isolated $CD150^{+}CD34^{-}c\text{-}Kit^{+}Sca1^{+}Lineage^{-}$ HSCs.

(f) Donor chimerism within the bone marrow and peripheral blood from freshly-isolated $CD150^{+}CD34^{-}c\text{-}Kit^{+}Sca1^{+}Lineage^{-}$ HSC KOs at 16-weeks post-transplantation. sgRNA 1 represented in black dots and sgRNA 2 represented in grey dots.

(g) Red blood cell (RBC) counts in transplant recipients from (H) at 16-weeks. Error bars represent s.e.m; P-value determined by one-way ANOVA.

(h) Platelet (Plt) counts in transplant recipients from (H) at 16-weeks. Error bars represent s.e.m; P-value determined by one-way ANOVA.

(i) Spleen weights of the transplant recipients from (H) at 16-weeks. Error bars represent s.e.m; P-value determined by one-way ANOVA.

**Figure S3: Analysis of SAGA component KO HSCs ex vivo**

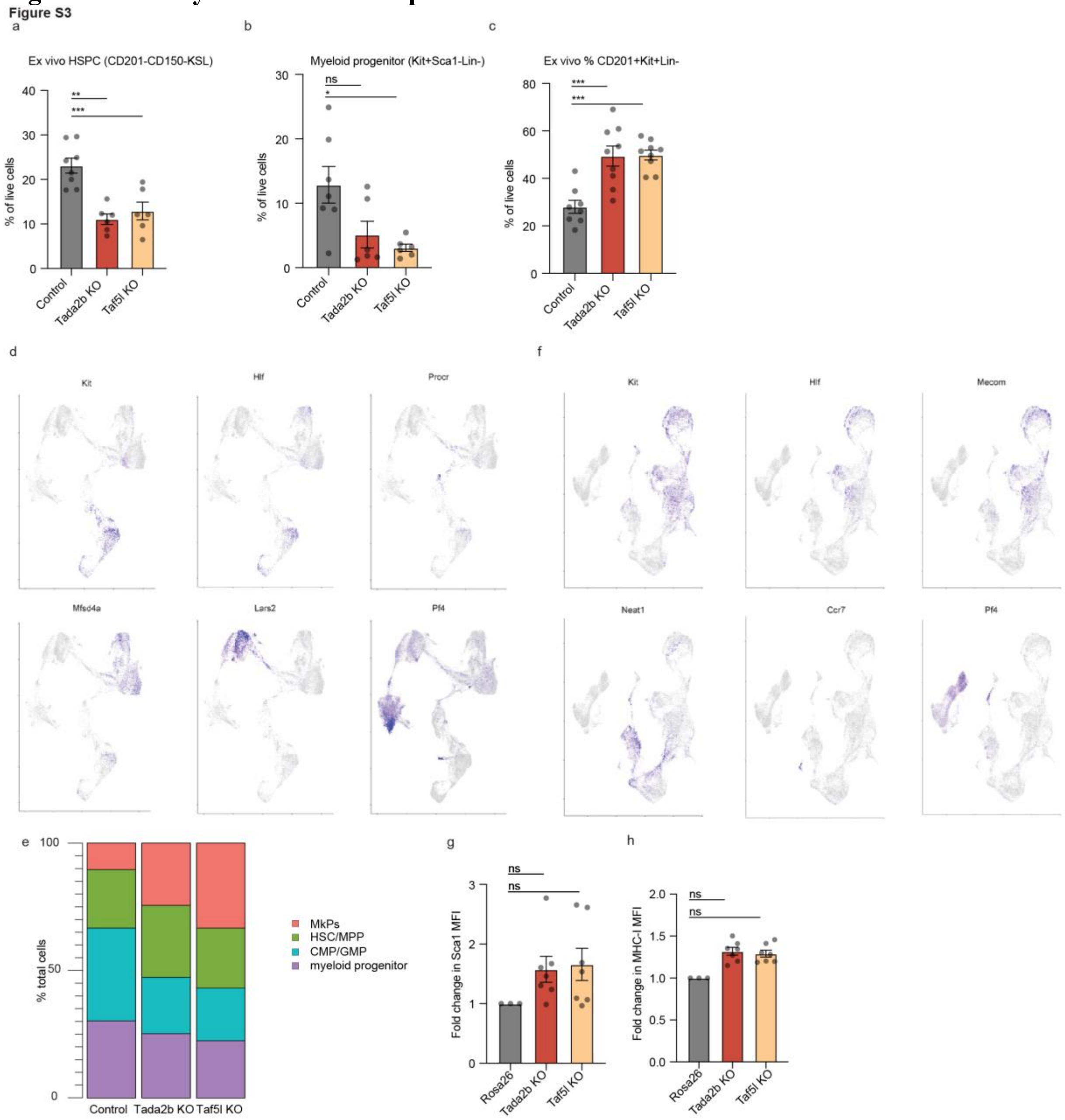

(a) Percentage of CD201$^-$CD150$^-$KSL progenitor cells in ex vivo cultures 14-days after RNP KO of SAGA complex members. Six biological replicates with two independent sgRNAs displayed. Error bars represent s.e.m; P-value determined by one-way ANOVA.

(b) Percentage of cKit$^+$Sca1$^-$Lineage$^-$ progenitor cells in ex vivo cultures 14-days after RNP KO of SAGA complex members. Six biological replicates with two independent sgRNAs displayed. Error bars represent s.e.m; P-value determined by one-way ANOVA.

(c) Percentage of CD201$^+$ cells in ex vivo cultures 14-days after RNP KO of SAGA complex members. Six biological replicates with two independent sgRNAs displayed. Error bars represent s.e.m; P-value determined by one-way ANOVA.

(d) Feature plots for genes defining clusters in complete single cell dataset.

(e) Feature plots for genes defining clusters in the HSC clusters.

(f) Frequency of cell type clusters in the control, *Tada2b* KO, and *Taf5l* KO cultures.

(g-h) Fold-change in Sca1 MFI (H) and MHC-I MFI (I) within ex vivo cultures 14-days after RNP KO of SAGA complex members. Six biological replicates with two independent sgRNAs displayed. Error bars represent s.e.m; P-value determined by one-way ANOVA.

### Figure S4: HSC transcriptional consequences of perturbing SAGA complex members

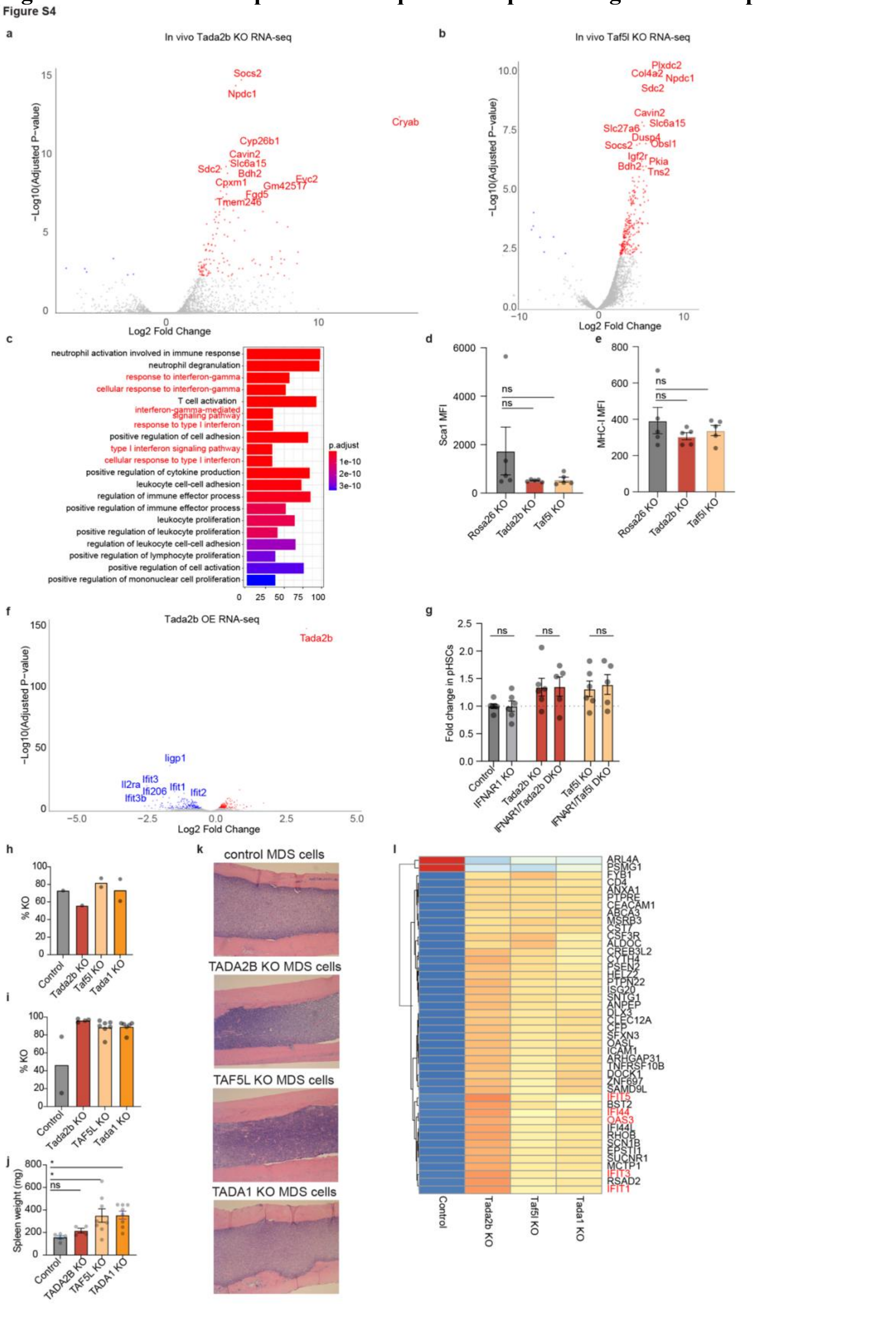

(a) Volcano plot comparing gene expression between in vivo control KSL HSPC with in vivo *Tada2b* KO KSL HSPCs, isolated at 16-weeks post-transplantation from recipient mice that received unexpanded RNP KO HSPCs (related to Figure S2f).

(b) Volcano plot comparing gene expression between in vivo control KSL HSPC with in vivo *Taf5l* KO KSL HSPCs, isolated at 16-weeks post-transplantation from recipient mice mice that received unexpanded RNP KO HSPCs (related to Figure S2f).

(c) GO enrichment of top 200 upregulated genes in Tada2b KO KSL HSPCs in vivo compared with control in vivo KSL HSPCs isolated at 16-weeks post-transplantation from recipient mice.

(d) Sca1 MFI within donor c-Kit$^+$Lineage$^-$ bone marrow cells at 12-weeks post transplantation in recipient mice described in Figure 1g-i.

(e) MHC-I MFI within donor c-Kit$^+$Lineage$^-$ bone marrow cells at 12-weeks post transplantation in recipient mice described in Figure 1g-i.

(f) Volcano plot comparing gene expression between GFP-transduced KSL HSCs and *Tada2b*-transduced KSL HSCs, at 3-days culture post-transduction.

(g) Fold change in phenotypic mouse HSCs (CD201$^+$CD150$^+$KSL) frequency within the indicated HSC cultures. Error bars represent s.e.m.

(h) Gene KO frequencies in human MDS-L cells prior to transplantation. One *TADA2B* sgRNA failed to induce any mutation and was excluded from the transplant.

(i) Gene KO frequencies in MDS-L sgRNA-transduced cells at 12-weeks post-transplantation, for mice described in Figure 4.

(j) Recipient mouse spleen weights at 12-week post-transplantation of control and KO MDS-L cells, for mice described in Figure 4.

(k) Representative H&E-stained bone marrow sections from recipient mice at 12-week post-transplantation, for mice described in Figure 4.

(l) Heatmap of gene expression within *in vivo* control and KO MDS cells.

**Materials Table: List of materials used in this study**

| REAGENT or RESOURCE | SOURCE | IDENTIFIER |
|---|---|---|
| Antibodies | | |
| Biotin anti-mouse CD4 | Biolegend | 100508 |
| Biotin anti-mouse CD8a | Biolegend | 100704 |
| Biotin anti-mouse TER-119/Erythroid Cells | Biolegend | 116204 |
| Biotin anti-mouse CD127 (IL-7Rα) | Biolegend | 135006 |
| Biotin anti-mouse Ly-6G/Ly-6C (Gr-1) | Biolegend | 108404 |
| Biotin anti-mouse/human CD45R/B220 | Biolegend | 103204 |
| APC/Cyanine7 Streptavidin | Biolegend | 405208 |
| FITC anti-mouse CD4 | Biolegend | 100510 |
| FITC anti-mouse CD8a | Biolegend | 100706 |
| FITC anti-mouse CD127 (IL-7Rα) | Biolegend | 135008 |
| FITC anti-mouse Ter119 | Biolegend | 116206 |
| FITC anti-mouse/human CD45R/B220 | Biolegend | 103206 |
| FITC anti-mouse Ly-6G/Ly-6C (Gr-1) | Biolegend | 108406 |
| Brilliant Violet 421 anti-mouse CD117 (c-Kit) | Biolegend | 105828 |
| PE anti-mouse Ly-6A/E (Sca-1) | Biolegend | 108108 |
| PE/Cyanine7 anti-mouse CD150 (SLAM) | Biolegend | 115914 |
| APC anti-mouse CD201 (EPCR) | Thermo Fisher | 17-2012-82 |
| PE/Cyanine7 anti-mouse CD45.1 | Biolegend | 110730 |
| eFluor 450 anti-mouse CD45.2 | Thermo Fisher | 48-0454-82 |
| FITC anti-mouse CD34 | Thermo Fisher | 11-0341-85 |
| Brilliant Violet 711 anti-mouse CD16/32 (Fcγ R III/II) | Biolegend | 101337 |
| APC anti-mouse CD117 (c-Kit) | Biolegend | 105812 |
| Alexa Fluor 700 anti-mouse H-2Kb (MHC class I) | Biolegend | 116522 |
| BUV395 Mouse Anti-Mouse CD45.2 | BD Biosciences | 564616 |
| Brilliant Violet 421 anti-mouse CD48 | Biolegend | 103428 |
| Brilliant Violet 785 anti-mouse CD150 (SLAM) | Biolegend | 115937 |
| APC anti-mouse CD4 | Biolegend | 100516 |
| APC anti-mouse CD8a | Biolegend | 100712 |
| PE anti-mouse/human CD11b | Biolegend | 101208 |
| PE anti-mouse Ly-6G/Ly-6C (Gr-1) | Biolegend | 108408 |
| APC/Cyanine7 anti-mouse/human CD45R/B220 | Biolegend | 103224 |
| Alexa Fluor 700 anti-mouse CD34 | Thermo Fisher | 56-0341-82 |
| APC anti-mouse Ter119 | Biolegend | 116212 |
| APC anti-mouse/human CD11b | Biolegend | 101212 |
| APC anti-mouse Ly-6G/Ly-6C (Gr-1) | Biolegend | 108412 |
| APC anti-mouse/human CD45R | Biolegend | 103212 |
| Microbead-conjugated anti-APC | Miltenyi | 130-090-855 |
| Microbead-conjugated anti-mouse CD117 (c-Kit) | Miltenyi | 130-097-146 |
| APC anti-human CD41 | Biolegend | 303710 |
| APC/Cy7 anti-human CD34 | Biolegend | 343514 |

| | | |
|---|---|---|
| PE anti-human CD201 | Biolegend | 351904 |
| BV785 anti-human CD45RA | Biolegend | 304140 |
| Anti-mouse Tada2b | St. John's Laboratory | STJ194361-200 |
| Anti-mouse Taf5l | Proteintech | 19274-AP |
| Anti-mouse Tada1 | Proteintech | 20337-1-AP |
| Anti-mouse histone H3 | Abcam | ab1791 |
| Anti-mouse Gapdh | Bethyl Laboratories | A300-641A |
| Peroxidase conjugated goat anti-rabbit IgG | Sigma Aldrich | A6667 |
| **Biological samples** | | |
| Umbilical cord blood CD34+ cells | NHSBT | NCB010 |
| **Chemicals, peptides, and recombinant proteins** | | |
| Propidium Iodide Solution | Sigma Aldrich | P4864 |
| Polyvinyl alcohol | Sigma Aldrich | P8136 |
| Soluplus | BASF | 50539897 |
| Recombinant Cas9 | IDT | 1081059 |
| Recombinant mouse thrombopoietin (THPO) | Peprotech | AF-315-14 |
| Recombinant mouse stem cell factor (SCF) | Peprotech | AF-250-03 |
| Recombinant human interleukin-3 (IL-3) | Peprotech | 200-03 |
| Recombinant human FLT3 ligand (FLT3L) | Peprotech | AF-300-19 |
| Recombinant mouse interferon alpha | Biolegend | 752804 |
| 740 Y-P | MedChemExpress | HY-P0175 |
| UM729 | Stem Cell Technologies | 72332 |
| Butzyamide | CellAid | N/A |
| ITSX | Thermo Fisher | 51500056 |
| PSG | Thermo Fisher | 10378016 |
| Ham's F12 media | Thermo Fisher | 11765054 |
| IMDM | Thermo Fisher | 12440061 |
| RPMI1640 | Thermo Fisher | 21875034 |
| Defined FBS | Corning | 35-010-CV |
| **Critical commercial assays** | | |
| Chromium GEM single cell 3' kit v3 | 10X Genomics | PN-1000268 |
| P3 Primary Cell Nucleofector Kit | Lonza | V4SP-3096 |
| Precision Plus Protein Kaleidoscope kit | Biorad | 1610375 |
| Quick-RNA microprep kit | Zymo Research | R1055 |
| SSIV Reverse Transcriptase kit | Thermo Fisher | 18090050 |
| **Experimental models: Cell lines** | | |
| NIH/3T3 cell line | ATCC | CRL-1658 |
| MDS-L cell line | Kida et al. 2018 | |
| **Organisms/strains** | | |
| B6J.129(Cg)-Gt(ROSA)26Sortm1.1(CAG-cas9*,-EGFP)Fezh/J | JAX | 026179 |
| PepboyJ | JAX | 002014 |

| | | |
|---|---|---|
| C57BL/6 | JAX | 000664 |
| C6.129S2-Ifnar1tm1Agt | MMRRC | 032045 |
| NOD.Cg-Prkdcscid Il2rgtm1SugTg(SV40/HTLV-IL3,CSF2)10-7Jic/JicTac | Taconics Biosciences | 13395-F |
| **Oligonucleotides** | | |
| Rosa26 sgRNA - ACUCCAGUCUUUCUAGAAGA | Thermo Fisher, Synthego | |
| Rosa26 fwd primer - ggctgttttggaggcaggaag | IDT | |
| Rosa26 rev primer - ccgaggcggatcacaagcaata | IDT | |
| Tada2b sgRNA 1 - CGGCGGACGAUUCACACUCU | Thermo Fisher, Synthego | |
| Tada2b sgRNA 2 - UGGGGUCCUGAGGCGGAGGG | Thermo Fisher, Synthego | |
| Tada2b sgRNA 3 - GCCGAAGCCGAACUGCUCGA | Thermo Fisher, Synthego | |
| Tada2b fwd primer - GGGATCTAGCTTGCTGCCAT | IDT | |
| Tada2b rev primer - GTACTGCTTGGCCGAGGTG | IDT | |
| Tada2b sequencing primer - ATAAGCCTGGCAACGCACAG | IDT | |
| Taf5l sgRNA 1 - UGGCACAACCAGAUUCUGAU | Thermo Fisher, Synthego | |
| Taf5l sgRNA 2 - UGGGGCUGCAGACACCGCAU | Thermo Fisher, Synthego | |
| Taf5l sgRNA 3 - AAACUGCACUUCAUACUGCU | Thermo Fisher, Synthego | |
| Taf5l fwd primer - ATTACGGTGCCAAAAGCCAC | IDT | |
| Taf5l rev primer - TCTCAGGAATATTGCGTGCT | IDT | |
| Taf5l sequencing primer - ATTGCGTGCTTCTCTTCTCT | IDT | |
| Taf6l sgRNA 1 - UUUUCCACUUGCAGUUCCAC | Thermo Fisher, Synthego | |
| Taf6l sgRNA 2 - CUCUCGUUCUGACAUGGCCC | Thermo Fisher, Synthego | |
| Taf6l sgRNA 3 - GGAGGUUUGUGGAGAUCCCU | Thermo Fisher, Synthego | |
| Taf6l fwd primer - TGCCCAGCGAGTGTTTTCTT | IDT | |
| Taf6l rev primer - CTTGGACTAGGGTCGTGCTG | IDT | |
| Taf6l sequencing primer - TTCTAGACACCTGACTGGCT | IDT | |

| | | |
|---|---|---|
| Tada1 sgRNA 1 - ACUGGGCCAACCUGAAGUUG | Thermo Fisher | |
| Tada1 sgRNA2 - GGCGACCUUUGUGAGCGAGC | Thermo Fisher | |
| Tada1 sgRNA 3 - UCUGCUUGAACCACAACUUC | Thermo Fisher | |
| Tada1 fwd primer (for sgRNAs 1 and 3) - GCTCATCTGAACGGAAGCGT | IDT | |
| Tada1 rev primer (for sgRNAs 1 and 3) - CACCCCCTTACCTGGTGTAG | IDT | |
| Tada1 sequencing primer (for sgRNAs 1 and 3) - ACCTCGTGGTCTCCTCCTAA | IDT | |
| Tada1 fwd primer (for sgRNA 2) - GCCGCGTTGATCTTTCGGTTGC | IDT | |
| Tada1 rev primer (for sgRNA 2) - ACGCCCAGGAGAAAGTCCTCCAG | IDT | |
| Mouse Ifit Taqman primers/probe | Thermo Fisher | Mm00515153_m1 |
| Mouse Gapdh Taqman primers/probe | Thermo Fisher | Mm99999915_g1 |
| **Recombinant DNA** | | |
| pLL3.7-shControl | Arede et al. 2022 | |
| pLL3.7-shTADA2B | Arede et al. 2022 | |
| pcDNA3.1-Ubc-GFP | This paper | |
| pcDNA3.1-Ubc-Tada2b-T2A-GFP | This paper | |
| pCMV-VSV-G-RSV-Rev | Addgene | 8454 |
| pMDLg/pRRE | Addgene | 12251 |
| pMD2.G | Addgene | 12259 |
| psPAX2 | Addgene | 12260 |
| Bassik mouse CRISPR knockout library | Addgene | 1000000121-1000000130 |
| pMCB306 | Addgene | 89360 |
| pLentiCas9-Blast | Addgene | 52962 |
| **Software** | | |
| FACSDiva | BD | Version 8 |
| FlowJo | BD | Version 10.8.0 |
| GraphPad Prism | GraphPad Software LLC | Version 10.1.1 |
| CasTLE | Morgens et al. 2016 | Version 1.0 |
| DESeq2 | Love et al. 2014 | Version 1.42.1 |
| Seurat | Butler et al. 2018 | Version 2.0 |
